# A new fold in TANGO1 evolved from SH3 domains for the export of bulky cargos

**DOI:** 10.1101/2022.02.02.478833

**Authors:** Oliver Arnolds, Raphael Stoll

## Abstract

Bulky cargos like procollagens, apolipoproteins, and mucins exceed the size of conventional COPII vesicles. During evolution a process emerged in metazoans, predominantly governed by the TANGO1 protein family, that organizes cargo at the exit sites of the endoplasmic reticulum and facilitates export by the formation of tunnel-like connections between the ER and Golgi. Cargo-recognition appears to be mediated by an SH3-like domain, however, just how the vastly different cargos are recognized remains elusive. Based on structural and dynamic data as well as interaction studies from NMR spectroscopy presented here, we show that the luminal cargo-recognition domain of TANGO1 adopts a new functional fold for which we suggest the term MOTH (MIA, Otoraplin, TALI/TANGO1 homology) domain. Also, differences in the structural properties within the domain family suggest fundamentally different mechanisms of cargo-recognition in vertebrates and invertebrates. Similarly, in vertebrates, it is proposed that cargo-specificity is mediated by structural differences within the vertebrate TANGO1 and TALI MOTH domains themselves.

## Introduction

The evolution of multicellular organisms (metazoa) brought forth the need to secret bulky cargos to supply the extracellular matrix with the building blocks it requires. ^1^ For this purpose, an intricate export machinery emerged that is predominantly governed by the transport and Golgi organization (TANGO) 1 protein family; these are transmembrane proteins located at the endoplasmic reticulum exit sites (ERES) found in most metazoans. ^2,3^ The simultaneous emergence of more complex organisms naturally led to a greater variety of large molecule-complexes that need to be exported. ^4^ Whereas only TANGO1 is present in invertebrates, vertebrates express different isoforms and a homologue termed TANGO1-like (TALI). ^2,5^

The TANGO1 protein family is responsible for organization of membranes and sorting of cargo at the ERES. The transmembrane proteins mediate the export of procollagens, apolipoprotein, and mucins, all of which exceed the size of conventional transport vesicles. ^5–7^ Formation of tunnel-like conduits between ERES and the Golgi apparatus enables export, facilitated by the cytosolic part of TANGO1 and TALI. ^2^

A domain annotated as SH3-like, present in the lumen of the endoplasmic reticulum of all known metazoan homologues of TANGO1, mediates cargo recognition and binding. ^3^ However, the process of cargo-recognition and binding remains elusive. The export of procollagen in vertebrates appears to depend on the formation of a ternary complex between procollagen, TANGO1’s cargo-recognition domain, and the vertebrate specific collagen-chaperone HSP47. ^6,8,9^ Contrarily, how procollagens are localized at the ERES in invertebrates remains unclear to date as they lack HSP47. ^2,10^ Furthermore, a mutation in TALI’s cargo-recognition domain leads to reduced secretion of apolipoproteins in mice, indicating the dependance of apolipoprotein secretion on the cargo-recognition domain. ^11^

This broad range of different cargos evokes the following questions: Are all vastly different types of cargo recognized by one single domain type or is cargo-specificity achieved in a less promiscuous way? With the luminal SH3-like domain of TANGO1 present in most metazoans, how do invertebrates export procollagen without HSP47? ^2,3,5,12^

SH3 domains are a family of non-catalytic protein-protein interaction modules located in the cytosol that are involved in a plethora of signaling pathways. ^13,14^ Typically, these domains adopt the highly conserved fold of a small β-barrel, which consists of five β-strands that form two perpendicular antiparallel β-sheets, three distinct loop regions (termed RT, nSrc, and distal), and a 3_10_-helix. ^13,15^ SH3 domains predominantly mediate protein-protein interactions by recognizing a left-handed polyproline-2 (PPII) helix. Two classes of consensus sequences, +xΦPxΦ P (class I) and ΦPxΦPx+ (class II) (with P for proline, Φ for a hydrophobic, + for a basic, and x for any residue), interact with a shallow hydrophobic region located between the nSrc and RT loop. ^13,16,17^

Our results presented here show that the characterization of the cargo-recognition domains of the TANGO1 protein family as SH3 or SH3-like domain might in fact be misleading, as it suggests a similar mode of operation (*a*) to SH3 domains and (*b*) within the domain family itself. We therefore propose that SH3 domains have evolved into a new functional fold present in TANGO1 to export bulky cargos in metazoans.

## Results

### TANGO1’s cargo-recognition domain adopts a modified SH3-like fold

In order to elucidate the molecular mechanisms that govern the cargo-recognition of the TANGO1 protein family we determined the structure of what hitherto has been termed as an SH3 domain using solution NMR spectroscopy. The sequence of the construct used corresponds to residues 21 to 131 of human TANGO1 (hsTANGO1(21-131)) and was based on the homology to the sequence of the MIA protein. Our high-precision structure ensemble exhibits backbone and heavy atom RMSDs over all secondary structure elements of 0.29±0.05 Å and 0.64±0.08 Å, respectively. (Supplementary Table 1) This structure revealed a typical small β-barrel fold, similar to SH3 domains, consisting of five antiparallel β-strands β2 (Y48-A52), β3 (D70-L78), β4 (V85-V90), β5 (T93-P98), and β6 (I102-E107). Additionally, a 3_10_-helix was identified between β-strands five and six, formed by residues K99 to L101. These elements, together with the three loop-regions (RT, nSrc, and distal), are also found in SH3 domains. ^13,16,17^ Notably, these features are extended by elongated termini that form two additional β-strands β1 (H35-C38) and β7 (L113-P116), which cover the classical SH3 fold in a lid-like manner. Moreover, the four cysteines form two disulfide bonds, thereby creating an additional loop (termed disulfide loop) between β-strands one and two, as well as tethering the unstructured C-terminus to the tip of the RT loop (Fig. 1a). These supplementary structural elements have already been observed in a similar fashion for the MIA protein ^18,19^ and are presumably present in other members of this domain family, i.e., TALI and Otoraplin, as judged from multiple sequence alignments (Fig. 1b). ^20^

**Figure 1:**
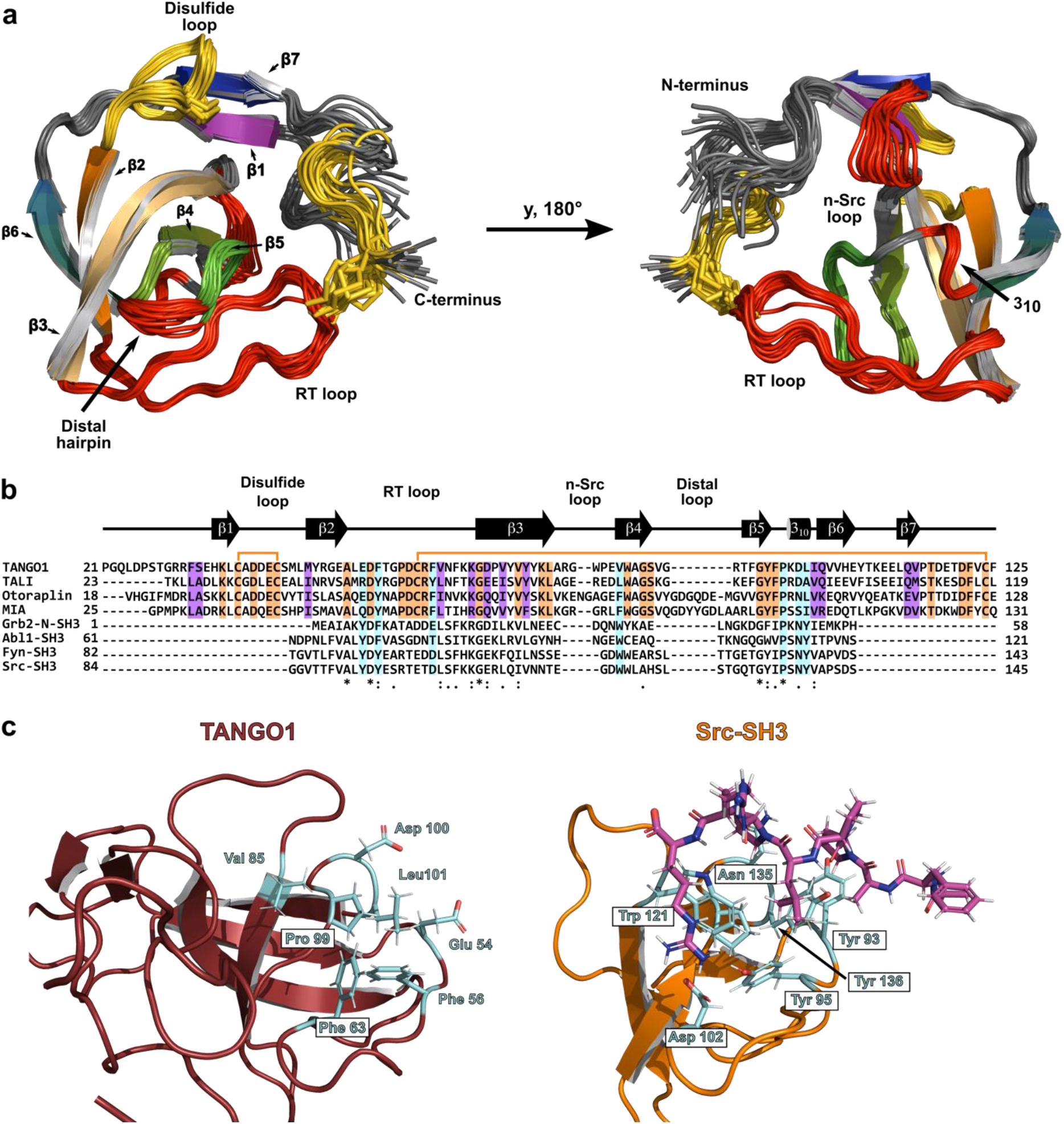
TANGO1’s cargo-recognition domain adopts a modified small β-barrel fold without retaining functional residues of SH3 domains. **a** Structure ensemble of human TANGO1’s cargo-recognition domain solved by solution NMR spectroscopy. Features also present in SH3 domains are highlighted in red, new features created by extended termini in yellow. Disulfide bridges are shown as sticks. (See also Supplementary Table 1) **b** Multiple sequence alignment of members of TANGO1 protein family and SH3 domains performed with ClustalOmega. Secondary structure elements of TANGO1’s domain are displayed above; disulfide bridges are represented by orange lines. Conserved and semi-conserved residues within the family are labelled in orange and purple, respectively. Residues, conserved within the domain family as well as SH3 domains are marked by an asterisk, semi-conserved residues by a colon. Residues that are critical for binding of PPII helices are highlighted in cyan, according to previous reports. ^17,22^ **c** Canonical binding site of PPII helices between the RT and nSrc loop on TANGO1’s domain (ruby) compared to the SH3 domain of human tyrosine-protein kinase Src (orange, PDB-ID: 1PRM). ^17^ Residues critical for binding are shown in cyan. The ligand peptide is displayed in magenta. (See also Supplementary Fig. 1 and 2)

### The canonical function of SH3 domains is abolished within the *mia* gene family

The TANGO1 protein family is encoded by genes of the *mia* family (*mia, mia2, mia3*, and *mial1*), all of which either consist of or carry a homologous domain described as an SH3 domain. ^20^ The eponymous MIA protein is an extracellular homologue to the cargo-recognition domains of TANGO1 and TALI, and has already been shown not to interact with classical PPII ligands of SH3 domains. ^19^ Due to the sequential and structural similarities of the cargo-recognition domains of the TANGO1 protein family with SH3 domains, we investigated their capability to interact with class I and II PPII helices that correspond to the recognition sequences of classical SH3 domains. The titration experiments using NMR spectroscopy analysis did not display any substantial chemical shifts perturbations (CSPs) of the backbone amide resonances of the domains from human TANGO1 and TALI as well as the MIA protein, indicating no interaction between the domains and peptides (Supplementary Fig. 1). Only Otoraplin exhibited CSPs at a high molar excess of a class II ligand, for which two-dimensional lineshape analysis revealed a low and physiologically probably irrelevant affinity with a dissociation constant K_D_ of 1.3±0.006 mM and a dissociation rate k_off_ of 3.5·10^5^±3.2·10^4^ s^-1^. However, CSP mapping to the surface of the predicted Otoraplin structure by DeepMind’s AlphaFold identified the interaction site to be at the disulfide and distal loop. ^21^ This is located on the opposite side of the protein compared to the interaction site of classical SH3 domains located between the RT and nSrc loops (Supplementary Fig. 2). Moreover, in case of Otoraplin, the interaction appears to be mainly driven by electrostatic forces between the negatively charged patch, comprised of the disulfide and distal loop, and the three arginine residues at the C-terminal end of the class II peptide. This might also explain why significant shifts were not observed for the class I ligand, which contains only a single arginine residue.

In spite of the structural homology to SH3 domains, many crucial residues required for binding of PPII helix ligands are not conserved in any of the four domains encoded by the *mia* gene family. ^17,22^ This abolishes the molecular basis for the interaction with PPII helix ligands, in good agreement with the results of the NMR-based titration experiments (Fig. 1b and c).

### Differences between the cargo-recognition domain in invertebrates and vertebrates

In vertebrates, procollagens are prepared for export at ERES by a ternary complex between TANGO1’s cargo-recognition domain, the vertebrate specific chaperone HSP47, and the procollagen itself. ^8,23^ In invertebrates, however, a conundrum emerges as they lack HSP47, yet the luminal cargo-recognition domain, previously annotated as an SH3, is apparently conserved on the domain level throughout metazoans. ^3,24^ Correspondingly, on a sequence level, stark differences between invertebrates and vertebrates can be observed (Fig. 2a). Both cysteines that form the highly conserved second disulfide bridge, thereby tethering the domain’s C-terminus to the RT loop in vertebrates, are absent in invertebrates (Fig. 2b). Using NMR spectroscopy, assignment of the backbone resonances of the sequence (Fig. 2a) from *Drosophila melanogaster* (dmTANGO1(30-139)) corresponding to hsTANGO1(21-131) provided first structural insights based on the chemical shift index (CSI). The chemical shift of NMR resonances depends on the chemical environment of the observed nucleus and therefore encodes structural information, from which the secondary structure element composition can be derived, revealed by the CSI. ^25^ This revealed a topology similar to the human domain, with two additional β-strands in the RT loop, which, however, are also observed for canonical SH3 domains. ^26^ Subsequent analysis of the pico-to nanosecond dynamics via the heteronuclear NOE (hetNOE) showed substantial differences in the dynamic properties (Fig. 2c and d). Firstly, both domains exhibit decreased hetNOEs for both termini, suggesting these to be unstructured, i.e., highly dynamic. However, for dmTANGO1(30-139) this is already observed for the region directly following β7. Conversely, the human domain only displays this behavior after the second disulfide bridge, i.e., the last cysteine (Fig. 2c, indicated in orange). Secondly, in the invertebrate domain the RT loop displays fast dynamics on the pico-to nanosecond timescale, which is completely rigid in the human domain. This is presumably due to the disulfide bridge that connects the RT loop to the C-terminus. In contrast, the nSrc loop and residues between β6 and β7 displayed decreased hetNOE values for the human domain, indicating fast dynamic structural fluctuations.

**Figure 2:**
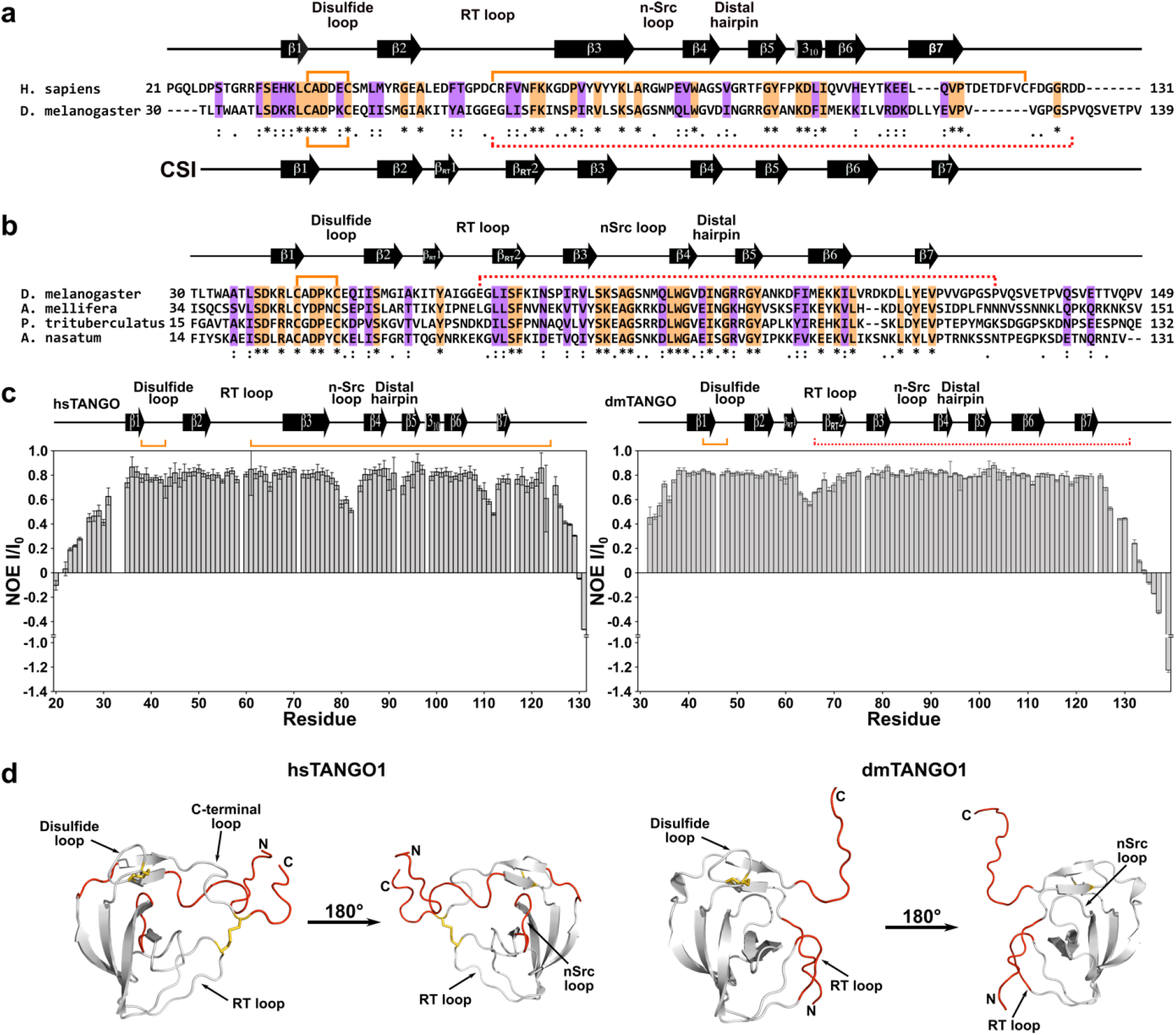
The cargo-recognition domain is conserved in invertebrates and distinctly different from the vertebrate domain. **a** Sequence alignment of TANGO1’s cargo-recognition domain from *Homo sapiens* and *Drosophila melanogaster* computed with ClustalOmega. Conserved and semi-conserved residues are displayed by asterisks and colons also highlighted in orange and purple, respectively. Disulfide bridges are indicated by orange lines, secondary structure elements above and below, according to the determined structure or the chemical shift index (CSI). The missing second disulfide bridge in invertebrates is indicated by dotted red line. (See also Supplementary Fig. 3) **b** Multiple sequence alignment of several invertebrate organisms. **c** hetNOE of TANGO1’s cargo-recognition domain from *Homo sapiens* and *Drosophila melanogaster*. Standard deviations calculated from all measurements (n=3) are displayed as error bars. **d** Regions that display motions on the pico-to nanosecond timescale projected to the structures of the domains from human and a predicted structure of *D. melanogaster* (AF-Q9VMA7-F1, accessed via https://alphafold.ebi.ac.uk/) in red. Disulfide bridges are shown as yellow sticks.

Finally, dmTANGO1(30-139) was tested for its interaction with PPII ligands as most residues in SH3 domains critical for the interaction are missing in invertebrate domains of TANGO1 as well. As for the human domain, an interaction between the domain and a PPII class II ligand could not be observed (Supplementary Fig. 3).

### Differences between the cargo-recognition domains of TANGO1 and TALI

In order to address the cargo-specificity of the TANGO1 protein family in vertebrates, we compared the structure of TANGO1’s domain with the structure predicted by AlphaFold for TALI’s domain from humans. Additionally, we compared our experimentally determined structure in solution of hsTANGO1(21-131) with the predicted structure by AlphaFold. Surprisingly, AlphaFold predicted residues 137 to 148, which were not included in the original construct for our structure determination, to form an amphipathic α-helix that is in contact with TANGO1’s core via hydrophobic residues between the RT and nSrc loop. Aromatic and hydrophobic residues located at this helix that are in contact with the interface at the RT loop appear to be conserved or at least semi-conserved in vertebrates (Fig. 3a). AlphaFold also predicts such a helix for mouse and zebrafish (Fig. 3b), though with varying degrees of confidence. Based on this prediction, a synthetic peptide that spans from residues 132 to 151 of human TANGO1 was titrated to the corresponding cargo-recognition domain using solution NMR spectroscopy. Subsequent CSP analysis revealed significant chemical shift differences of the amide resonances surrounding the RT and nSrc loop as well as the 3_10_-helix, in good agreement with the predicted structure (Fig. 3c and d). Residues displaying shift differences exceeding twice the standard deviation and with a relative surface accessibility of 30 % or more were used for two-dimensional lineshape analysis, yielding a dissociation constant of 320.8±4.1 μM with a dissociation rate k_off_ of 1.1·10^3^±29.1os^-1^ (Supplementary Fig. 4c).

**Figure 3:**
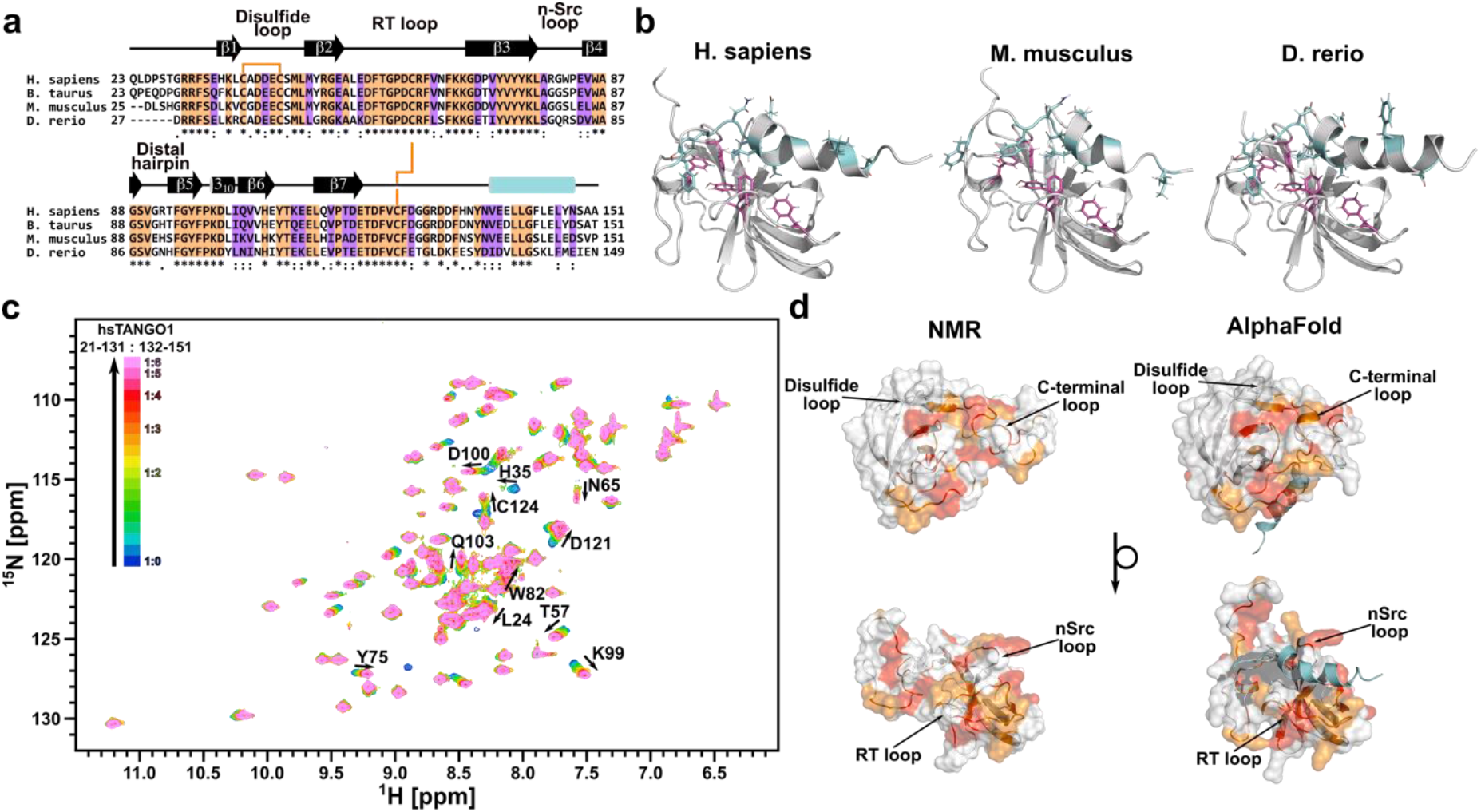
The predicted C-terminal helix is conserved and binds to TANGO1’s cargo-recognition domain. **a** Multiple sequence alignment of the cargo-recognition domain of TANGO1 from different vertebrate organisms. Conserved and semi-conserved residues are highlighted in orange and purple, respectively. Conserved disulfide bridges are indicated by orange lines. **b** TANGO1’s cargo-recognition domain displays a C-terminal helix that is conserved throughout vertebrate organisms, e.g., human, mouse, and zebrafish (AF-Q5JRA6-F1, AF-Q8BI84-F1, and AF-F1R5N2-F1, respectively. Accessed via https://alphafold.ebi.ac.uk/). Conserved aromatic residues within the domain are shown as magenta, conserved residues corresponding to the region of the synthetic peptide as cyan sticks. **c** 2D ^1^H ^15^N-HSQC titration spectra. Labelled residues were used for subsequent determination of K_D_ and k_off_ values. (See also Supplementary Fig. 4) **d** Residues from CSP analysis exceeding single and double standard deviations based on the average shift differences for all residues projected onto the surface structure of TANGO1’s cargo-recognition domain. Experimental NMR structure on the left, AlphaFold prediction including C-terminal helix (cyan) on the right.

This C-terminal helix appears to be conserved for TANGO1 throughout vertebrates, which, however, is not the case for TALI (Fig. 4a and b). Instead, residues located C-terminally to the second disulfide bridge in TALI seem to be unstructured and not conserved.

**Figure 4:**
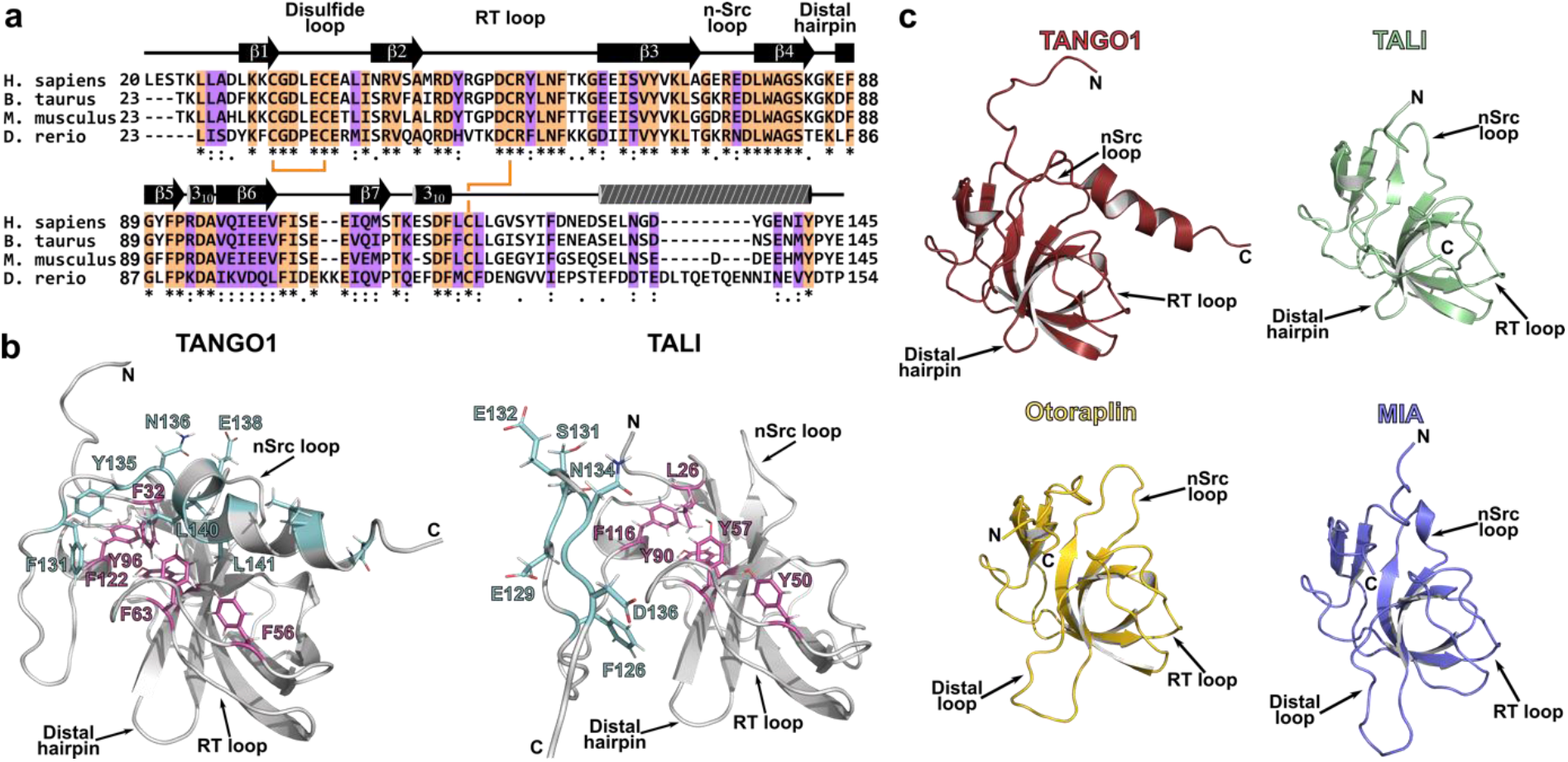
A conserved C-terminal helix is unique for vertebrate TANGO1. **a** Multiple sequence alignment of the cargo-recognition domain of TALI from different vertebrate organisms. Conserved and semi-conserved residues are highlighted in orange and purple, respectively. Conserved disulfide bridges are indicated by orange lines. C-terminal helix found in TANGO1 is indicated by striped cylinder. **b** Structures of human TANGO1’s and TALI’s cargo-recognition domain predicted by AlphaFold. Conserved aromatic residues within the domain are shown as magenta, conserved and semi-conserved residues corresponding to the region of the helix in TANGO1 as cyan sticks. **c** Domain family encoded by the *mia* gene family. Structures for TANGO1, TALI, and Otoraplin are AlphaFold predictions (AF-Q5JRA6-F1, AF-Q96PC5-F1, and AF-Q9NRC9-F1, respectively. Accessed via https://alphafold.ebi.ac.uk/). The RCSB-PDB database entry for MIA is 1I1J. ^18^

## Discussion

Based on sequence homology, the luminal cargo-recognition domain of TANGO1 has previously been annotated as an SH3 domain. Notably, the UniProt database assigned the SH3 domain to residues 45 to 107 (UniProt entry Q5JRA6) but neglected terminal extensions that are conserved throughout the *mia* gene family. Indeed, the structure presented here revealed a small β-barrel fold at the domain’s core, which is, importantly, complemented by terminal elongations. These create two additional β-strands and two disulfide bridges that tether these new features to the classical SH3 fold (Fig. 1a). Whereas previous reports already showed similar structural features for the MIA protein, a sequence alignment and structures predicted by AlphaFold for TALI’s cargo-recognition domain and Otoraplin strongly suggest that this is indeed the case for all members of this domain family (Fig. 4c). ^18,19^ Furthermore, we report that none of these domains retained the ability to interact with PPII helix motifs in a manner that SH3 domains do. The interaction of SH3 domains with PPII ligands is classically driven by conserved aromatic and hydrophobic residues, most of which are not conserved in all members of the *mia* gene family (Fig. 1b). ^13,16,17^ Hence, we propose that the stable fold of the small β-barrel has been adapted and modified for new physiological tasks in non-cytosolic space. During this evolutionary process, the canonical SH3 function has not been retained, which has been observed for Sm-like domains in a similar fashion. ^27^ This motivates us to suggest a new name for this domain family in order to distinguish it from SH3 domains: the MOTH (MIA, Otoraplin, TALI/TANGO1 homology) domain.

As we show here, an evolutionary intermediate between SH3 domains and the vertebrate MOTH domains can already be observed in invertebrates, such as *D. melanogaster*, which has also lost the ability to interact with PPII helix ligands (Supplementary Fig. 3). Whereas the extended termini and first disulfide bridge have already emerged to adopt a similar topology present in the MOTH domains as indicated by the CSI (Fig. 2a), the second disulfide bridge is notably missing. Dynamic data from heteronuclear NOE NMR experiments show that this leaves the residues C-terminal of the last β-strand (β7) completely unstructured, as the C-terminus is not tethered to the RT loop. Furthermore, different regions of hsTANGO1(21-131) and dmTANGO1(30-139) display pronounced dynamic properties (Fig. 2c, d). dmTANGO1(30-139) appears to be rather rigid, with only the RT loop exhibiting movements on the pico-to nanosecond timescale, which is probably facilitated by the missing disulfide bridge. Conversely, these regions are rather rigidified in hsTANGO1, while the nSrc loop and the unstructured part between β6 and β7 were found to be flexible. Due to these structural and dynamic differences, we hypothesize that the mechanism by which bulky cargo is recognized in invertebrates is fundamentally different from the underlying processes present in vertebrates. Interestingly, the amino acid composition for the domain in both phyla varies to an extent that leads to drastically different isoelectrical points - 4.64 for hsTANGO1(21-131) and 8.97 for dmTANGO1(30-139) - calculated by Expasy’s ProtParam tool. ^28^ This coincides with the observation that two homologous domains emerged in invertebrates and vertebrates that are, nonetheless, conserved within the respected phylum (Fig. 2b and 3a). We further suggest that the more complex catalogue of bulky cargo in vertebrates, especially illustrated by the greater variety of collagens, was facilitated by the concomitant changes to the cargo-recognition domain and also by the emergence of novel molecular mechanisms. The interaction with the chaperone HSP47, which in turn recognizes a plethora of different procollagens itself, clearly serves as a primary example in this context.

In vertebrates, the MOTH domains of TANGO1 and TALI have been observed to mediate the export of bulky cargo, thereby raising questions about the cargo-specificity of these domains. Here, we report a hitherto undiscovered α-helix C-terminal of TANGO1’s MOTH domain that appears to be conserved in vertebrates (Fig. 3a, b), based on structures predicted by AlphaFold. NMR-based titration experiments with a peptide corresponding to the residues forming the α-helix in TANGO1’s MOTH domain indicate binding of the helix at the predicted interaction site. Due to its conservation throughout different vertebrate organisms, we propose this motif to be of functional significance, as no other MOTH domain contains this helix (Fig. 4c). This is corroborated by the structure presented here for TANGO1’s MOTH domain that lacks the helix but still adopts a stable fold, implying no or very little structural relevance (Fig. 1a). Notably, this α-helix appears to be non-essential for the interaction between HSP47 and TANGO1’s MOTH domain, as previous reports on the interaction with HSP47 used a shortened construct of the MOTH domain that did not contain the residues forming this C-terminal helix. ^8^ Therefore, we hypothesize either a modulating role or a yet to be discovered (secondary) interaction for TANGO1’s MOTH domain mediated by this C-terminal helix. Ultimately, these structural differences seen in TALI and TANGO1 infer that cargo-specificity for the export of bulky cargo may be mediated by the respective MOTH domain, in accordance with both being expressed in different tissues. ^5,29,30^ Our results presented here shed light on the foundation of the cargo-recognition of bulky molecules and may ultimately aid drug development for diseases like fibrosis in which regulation of cargo export is impaired.

## Methods

### Constructs

MOTH domains of human TALI (23-123), Otoraplin (18-128), and MIA (19-131), and TANGO1 (30-139) from *Drosophila melanogaster* (dmTANGO1(30-139)) were expressed from codon-optimized sequences in a modified pQE40 expression vector ^31^ in M15 pRep4 *E. coli* strain. Human TANGO1 (21-131) (hsTANGO1(21-131)) was expressed from codon-optimized sequences in a modified pET19b vector gifted from Matthias Lübben, PhD of the department of Biophysics at the Ruhr University of Bochum in BL21(DE3)RIL *E. coli* strain. The sequences of the used constructs were based on the homology to the sequence of the MIA protein.

### Protein expression

Cells were typically grown in custom minimal medium (0,3 mM CaCl_2_, 1 mM MgCl_2_, 3 ml/l 100x BME vitamins, 50.0 mg/l EDTA, 8.3 mg/l FeCl3 x 6 H_2_O, 0.84 mg/l ZnCl_2_, 0.13 mg/l CuCl_2_ x 2 H_2_O, 0.1 mg/l CoCl_2_ x 6 H_2_O, 0.1 mg/l boric acid, 13.5 μg/l MnCl_2_ x 4 H_2_O, 10 g/l ^12^C-D-glucose, 5 ml/l ^12^C-glycerol, 2 g ^14^N-NH_4_Cl, 42,3 mM Na_2_HPO_4_ x 2 H_2_O, and 22 mM KH_2_PO_4_; pH adjusted to 7.4). For expression, cells were incubated in isotopically enriched medium. For this ^13^C6-D-glucose (4 g/l) and ^15^N-NH_4_Cl (2 g/l) were used to substitute their respective isotopes. Crucially, no glycerol was added to media, if ^13^C-enrichment was required.

Chemically competent cells were transformed with respective plasmid DNA and then transferred to 200 ml of minimal medium, which was incubated overnight at 37 °C. 2 l of minimal medium were inoculated from the pre-culture to an OD_600nm_ of 0.1. The culture was incubated at 37 °C and to an OD_600nm_ of 0.8 – 1.0. Next, the cells were harvested by centrifugation at 37 °C and 3,000 x *g* for 10 min. The resulting pellets were re-suspended in 500 ml isotopically enriched medium. Expression was induced directly after re-suspension by 1 mM IPTG. The culture was incubated for 21 h at 30 °C. Cells were harvested by centrifugation at 4 °C and 3,000 x *g* for 10 min and resulting pellets re-suspended in 50 mM Tris/HCl, 1 mM EDTA, pH 8.

### Protein purification from inclusion bodies

Cells were mechanically lysed by micro-fluidization. The resulting homogenates were cleared by centrifugation at 10,000 x *g* and 20 °C for 30 min. Pelleted inclusion bodies were re-suspended in 50 mM Tris/HCl, 1 mM EDTA, 1 % Triton X-100, pH 8 by vortexing vigorously for 5 min. The suspension was cleared again by centrifugation at 7,500 x *g* and 20 °C for 10 min and the supernatant discarded. This was repeated until the supernatant was clear. Afterwards, the pellet was re-suspended in 50 mM Tris/HCl, 1 mM EDTA, pH 8 by vortexing and the suspension again cleared by centrifugation. This process was repeated until no more detergent was observed. Inclusion bodies were solubilized at room temperature in 15 ml of 6 M guanidinium chloride, 12.5 mM NaHCO_3_, 87.5 mM Na_2_CO_3_, 0.2 M DTT, pH 10. Following this, the pH of the solution was adjusted to 3 and then cleared by centrifugation at 10,000 x *g* and 20 °C for 30 min. The cleared opaque supernatant was filtered using a Filtropur S 0.45 μM filter (Sarstedt). Buffer of the filtered solution was then exchanged to 3 M guanidinium chloride, 4.7 mM sodium citrate dihydrate, 45.7 mM citric acid, pH 3. Afterwards, the DTT-free solution was then dropped very slowly under stirring to refolding buffer (1 M arginine hydrochloride, 50 mM Tris/HCl, 1 mM EDTA, pH 8 and different ratios of oxidized and reduced glutathione, depending on the protein).

Solubilized inclusion bodies were diluted 1:200 in refolding buffer with 0.5 mM of oxidized and 2.5 mM reduced glutathione and incubated at room temperature for 3 days. Subsequently, 2.5 volumes of 50 mM Tris/HCl, 1 mM EDTA, pH8 were added to the solution and filtered through folded paper filters. Then, the volume was reduced to a volume of 400 ml using the *ÄktaFlux* system with a 3 kDa MWCO cartridge (GE Healthcare). Afterwards the system was used for buffer exchange with 50 mM HEPES, pH 8, and 1 mM EDTA. The N-terminal His-tag was subsequently cleaved off with 0.6 mg of TEV-protease by incubating for 17 h at 20 °C. TEV-protease and unprocessed hsTANGO1 were removed by Protino-Kit (Macherey-Nagel). Flowthrough and wash fractions were collected, and monomeric protein was purified by size-exclusion chromatography using a HiLoad™ 26/600 Superdex™ 75 pg (Merck) equilibrated with 25 mM HEPES, 150 mM NaCl and pH 7.4.

Redox ratios during refolding of oxidized:reduced glutathione were 0.5 mM:2.5 mM for dmTANGO1(30-139), 0.5 mM:0.5 mM for TALI(23-123) and Otoraplin, and 0.5 mM:5.0 mM for MIA. Otoraplin and MIA were incubated with a 1:200 dilution at room temperature for 3 days, dmTANGO1(30-139) with 1:200 at 8 °C for 3 days, and TALI(23-123) with 1:40 at 8 °C for 1,5 days. Purification of Otorpalin’s and TALI(23-123)’s MOTH domain as well as dmTANGO1(30-139)’s cargo-recognition domain was done as described for hsTANGO1(21-131). PBS buffer at pH 7,4 was used for buffer exchange with the *ÄktaFlux* system as well as for equilibration for size-exclusion chromatography.

After refolding MIA’s MOTH domain, 1.4 M ammonium sulfate was added and then applied to a hydrophobic interaction chromatography column with a Toyopearl Butyl-650S-substituted matrix (Tosoh Bioscience) using a peristaltic pump. After washing with 50 ml of 50 mM Tris/HCl, pH 7.4, and 1.4 M (NH4)2SO4, the protein was eluted by a step-gradient with a decreasing concentration of ammonium sulfate by 0.2 M per 50 ml step to a final concentration of 0 M. Eluted fractions were analyzed by SDS-PAGE, fractions containing protein were pooled together, and combined fractions were dialyzed twice with a 3 kDa MWCO membrane in 5 l of PBS buffer, pH 7.4. After concentrating, size-exclusion chromatography equilibrated with PBS buffer at pH 7.4 was used to extract monomeric protein. (adapted from ^31^)

Protein concentrations were determined for hsTANGO1(21-131), dmTANGO1(30-139), FDP, and MIA via the absorbance at 280 nm and the Lambert-Beer law using molar attenuation coefficients predicted by ExPASy’s ProtParam webtool. ^28,32^ Concentration of TALI(23-123)’s MOTH domain was determined using the Pierce™ BCA Protein Assay Kit (Thermo Scientific).

All monomeric protein solutions were further concentrated, aliquoted, snap-frozen in liquid nitrogen and stored at −80 °C.

### Solution NMR spectroscopy

All spectra (see Supplementary Table 2) were recorded on Bruker DRX 600 and AVANCE III HD 700 spectrometers at 298 K. Typically, samples of 1 mM [U-^15^N-^13^C]-enriched protein were measured for three- or four-dimensional spectra. For titration analysis, [U-^15^N]-enriched protein samples of 0.2 mM were prepared, which is described in detail below. Interscan delay was typically set to 1 s. Mixing time for NOESY experiments was set to 120 ms. Spectra were referenced to the methyl signal of DSS, processed with Topspin 3.6.1, and subsequently assigned and analyzed using CcpNmr Analysis 2.4.2. ^33^ MOTH domain of hsTANGO1(21-131) was typically measured in 25 mM HEPES, pH 7.4, 150 mM NaCl, 1 % CHAPS, 10 % D_2_O, 0.02 % (w/v) NaN_3_, and DSS. All other MOTH domains and dmTANGO1(30-139) were measured in PBS buffer, pH 7.4, 10 % D_2_O, 0.02 % (w/v) NaN_3_, and DSS.

For the assignment of backbone and side chain resonances of hsTANGO1(21-131), threedimensional HNCO, HNcaCO, HNCA, CBCAcoNH, HNCACB, hCCcoNH, HcccoNH, and ^1^H^15^N^1^H-NOESY as well as four-dimensional ^1^H^15^N^1^H^13^C-NOESY spectra were recorded. For threedimensional HCCH-COSY, HCCH-TOCSY, and ^1^H^13^C^1^H-NOESY as well as four-dimensional ^1^H^13^C^1^H^13^C-NOESY that do not require an amide proton for detection, the sample was lyophilized and re-solvated in 100 % D_2_O. For the assignment of the backbone resonances of TANGO1’s cargo-recognition domain from *D. melanogaster*, HNCO, HNcaCO, HNCA, HNcoCACB, and HNCACB spectra were recorded. Backbone resonances from *D. melanogaster* were analyzed using the webtool CSI 3.0 to extract structural information in form of the chemical shift index. ^34^

For titration experiments, two-dimensional ^1^H^15^N-HSQC spectra were recorded for are reference containing only protein and subsequently after adding a small amount of peptide from a high-concentrated stock solution. All synthetic peptides were resolved in the buffer used for the respective protein and the pH adjusted to 7.4.

For the interaction of a class I PPII helix peptide, spectra of hsTANGO1(21-131) (215 μM), TALI(23-123) (196 μM), Otoraplin (165 μM), MIA (182 μM), and dmTANGO1(30-139) (160 μM) were recorded with a 10-fold molar excess of from a stock solution of 16.5 mM p85α(91-104), each. The interaction with a class II PPII helix peptide (SOS1 (1149-1158)) was investigated by a 10-fold molar excess of peptide to protein for hsTANGO1(21-131) (193 μM), TALI(23-123) (200 μM), and Otoraplin (168 μM), a 24-fold excess was used for MIA (200 μM) from a 20 mM stock solution. To determine the dissociation constant of SOS1 (1149-1158) and Otoraplin, a series of titration spectra with increasing amounts of peptide were recorded with molar ratios of protein to peptide of 1:0.5, 1.0, 1.5, 2.0, 2.5, 3.0, 3.5, 4.0, 4.5, 5.0, 6.0, 7.0, 10.0, 15.0, 20.0, 25.0, 30.0, 40.0, and 50.0.

For hsTANGO1(21-131)’s interaction with the synthetic peptide corresponding to residues 132-151, a sample of 208 μM hsTANGO1(21-131) in 25 mM HEPES, pH 7.4, 150 mM NaCl, 10 % D_2_O, 0.02 % (w/v) NaN_3_, and DSS was measured as reference. Peptide was added corresponding to molar ratios of 1:0.25, 0.5, 0.75, 1.0, 1.25, 1.5, 1.75, 2.0, 2.25, 2.5, 2.75, 3.0, 3.25, 3.5, 4.0, 5.0, and 6.0 from a stock solution of 24.1 mM.

Heteronuclear NOE spectra were recorded as pseudo-three-dimensional spectra on samples of 1 mM [U-^15^N]-enriched protein as triplicates with an interscan delay of 5 s. Signals were picked in CcpNMR Analysis 2.4.2.

### Structure calculation

Distance restraints were extracted from the initial peak lists of all NOESY spectra after complete side chain assignment. Dihedral restraints based on predicted Φ - and Ψ-torsion angles by TALOS+, disulfide bridges between cysteines 38 and 43 as well as 61 and 124, and the *cis*-conformation of proline 83 were set as additional restraints. ^35^ Structures were calculated using a two-step-approach in ARIA 2.3.1. ^36^ In both steps, the algorithm for torsion angle dynamics was applied for the simulated annealing protocol of the molecular dynamics simulation. Folded conformations were computed from an unstructured, extended strand. The total energy of a calculated structure was used as a sort criterion for the resulting coordinate files. First, ARIA 2.3.1 was utilized to complete the assignment of interresidual proton-proton-contacts using nine iterations with decreasing violation thresholds and number of calculated structures (see Supplementary Table 3). For the assignment process itself, network anchoring and for the distance restraint potential a log-harmonic shape was used, which included Bayesian weighting of the distant restraints in order to increase the quality of assignments. After completion of a full calculation protocol, all distance violations >0.5 Å were systematically checked and reassigned, if necessary. Dihedral restraint violations between 5 ° and 12 ° were excluded from the calculation, because the uncertainty of the Φ- and Ψ-torsion angles predicted by TALOS+ was reported as 12.6 ° and 12.3 °, respectively. ^35^ In the second step, a structural ensemble was calculated from a fully assigned peak list and subsequently refined in explicit water. ^37^ Because the log-harmonic potential was not compatible with the refinement in explicit solvent, the more traditional flat-bottom potential shape was used in this second approach. For this, a structural ensemble was calculated from an extended strand as a starting point in ARIA 2.3.1 as well, but only a single iteration was applied. An initial structure ensemble from previous calculated structures was used as a reference for the assignment step of ambiguous restraints during the ARIA protocol. 200 structures with the lowest total energy were refined in explicit water. From these, 20 structures without any NOE or dihedral violations were chosen for the final ensemble based on the lowest values in three energy terms with decreasing priority (i.e., total energy, NOE energy, and van-der-Waals energy). Finally, structure quality analysis of this last ensemble was carried out by PROCHECK-NMR. ^38,39^ Assignment of secondary structure elements displayed in this paper is based on the analysis of the STRIDE webserver. ^40,41^

### Chemical shift perturbation analysis

In order to determine the binding site and dissociation constant, only residues with a shift difference exceeding twice the standard deviation (SD) and displaying a relative surface accessibility of at least 30 % according to PyMOL were used for further analysis. ^42^ Determination of the dissociation constant via TITAN required processing of the spectra using nmrPipe. ^43,44^ Two-dimensional ^1^H-^15^N HSQC spectra were processed with an exponential window function for apodization with 4 Hz and 10 Hz of exponential line broadening in the proton and nitrogen dimension, respectively. Signals of residues meeting the aforementioned criteria were fitted with a simple two-state binding model with subsequent bootstrap error analysis.

### Multiple sequence alignment

Multiple sequence alignments were done with ClustalOmega from the EBI tools webservice. ^45,46^ Amino acid sequences were taken from UniProt database (https://www.uniprot.org/) using entries for vertebrate TANGO1 from *Homo sapiens* (Q5JRA6), *Bos taurus* (Q0VC16), *Mus musculus* (Q8BI84), and *Danio rerio* (F1R5N2). Invertebrate sequences for TANGO1 were used from *Drosophila melanogaster* (Q9VMA7), *Portunus trituberculatus* (A0A5B7CJZ6), and *Armadillidium nasatum* (A0A5N5SML6). Entries for TALI were used from *Homo sapiens* (Q96PC5), *Bos taurus* (A0A3Q1LM15), *Mus musculus* (Q91ZV0), and *Danio rerio* (A5PLB3). Human sequences for Otoraplin and MIA correspond to database entries Q9NRC9 and Q16674, respectively. Exemplary sequences for SH3 domains were used from human proto-oncogene tyrosine-protein kinase Src (P12931), growth factor receptor-bound protein 2 (P62993), tyrosine-protein kinase ABL1 (P00519), and tyrosine-protein kinase Fyn (P06241).

TANGO1 sequence from *Apis mellifera* was taken from NCBI’s gene database (https://www.ncbi.nlm.nih.gov/) entry LOC412103.

### Quantification and statistical analysis

Standard deviations (SD) for all CSP analyses (Fig. 3 and Supplementary Fig. 2 and 4) were calculated based on the average shift differences observed for all signals of the ^1^H^15^N-HSQC spectra with Microsoft Excel for Mac (v16.43). Heteronuclear NOEs (Fig. 2) were quantified by forming the ratio of peak heights of the saturated spectra and the non-saturated reference spectra. Displayed are averaged values (n=3), error bars indicated standard deviation of these averaged values calculated by CcpNMR Analysis 2.4.2. Dissociation constants K_D_ and rates k_off_ were determined by an iterative fitting procedure to a two-state binding model using with subsequent bootstrap error analysis implemented in TITAN (Supplementary Fig. 4). ^44^

## Data availability

The NMR assignments for TANGO1(30-139) from *Drosophila melanogaster* and human MOTH domain of TANGO1(21-131) are deposited in the Biological Magnetic Resonance Bank (https://bmrb.io/) under accession codes XXX and XXX, respectively. The atomic coordinates for the structure in solution of human TANGO1(21-131) have been deposited with the Protein Data Bank (https://www.rcsb.org/) under accession code XXX.

## Acknowledgements

We thank Stefanie Pütz for expert technical assistance and Xueyin Zhong for stimulating discussions. O.A. thanks the RUB Research School^Plus^ for financial support. R.S. gratefully acknowledges support from the DFG (grant nos. INST213/757-1 FUGG and INST 213/843-1 FUGG).

## Author contributions

O.A. and R.S. conceived the research, prepared the figures, and wrote the manuscript. O.A. performed all experiments. O.A. and R.S. recorded and analyzed the NMR data.

## Additional information

**Correspondence** and requests for materials should be addressed to Raphael Stoll.

