## Supplemental Information for "A new fold in TANGO1 evolved from SH3 domains for the export of bulky cargos"

### Supplementary Table 1.

**Statistics for all conformational restraints used in the calculation and geometric quality statistics for the final NMR ensemble of the 20 lowest energy structures for TANGO1's cargo-recognition domain.** Deviations ± standard deviations of this ensemble from averaged coordinates are summarized in the RMSD.

| **Conformational restraints** | |
|---|---|
| Distance restraints | |
| Intraresidual (i = j) | 815 |
| Sequential (\|i - j\| = 1) | 314 |
| Short-range (2 < \|i - j\| < 3) | 69 |
| Medium-range (4 < \|i - j\| < 5) | 27 |
| Long-range (\|i - j\| ≥ 5) | 404 |
| Ambiguous | 175 |
| Dihedral restraints (Φ/Ψ) | 195 |
| Other structural restraints | |
| Oxidized cysteines | 4 |
| *cis*-prolines | 1 |
| **Total number of restraints** | **2004** |

| **Structure quality** | |
|---|---|
| Average RMSD of secondary structures | [Å] |
| Backbone | 0.29 ± 0.05 |
| Heavy atoms | 0.64 ± 0.08 |
| Ramachandran statistics | [%residues] |
| Core regions | 80.1 |
| Allowed regions | 18.3 |
| Generous regions | 0.7 |
| Disallowed regions | 1.0 |

### Supplementary Table 2.

#### Pulse programs used for NMR spectroscopy.

Pulse programs and parameters such as number of scans (NS), amount of recorded data points using non-uniform sampling (NUS), sweep width (SW), and points recorded in the time domain (TD) used for NMR spectroscopy.

| Pulse program | | | F1 | | F2 | | F3 | | F4 | | Ref |
|---|---|---|---|---|---|---|---|---|---|---|---|
| | NS | NUS | SW [ppm] | TD | SW [ppm] | TD | SW [ppm] | TD | SW [ppm] | TD | |
| zggpw5 | 8 | - | 15.9 | 32768 | - | - | - | - | - | - | [1] |
| hsqcfpf3gpphwg | 8 | - | 29.4 | 256 | 15.9 | 2048 | - | - | - | - | [2–5] |
| hsqcctetgpsp | 32 | - | 80.0 | 256 | 13.0 | 1024 | - | - | - | - | [6] |
| hncagpwg3d | 16 | - | 32.0 | 128 | 29.4 | 48 | 15.9 | 2048 | - | - | [7–9] |
| hncacbgpwg3d | 128 | 30% | 80 | 128 | 29.4 | 40 | 15.9 | 2048 | - | - | [10,11] |
| hncogpwg3d | 16 | - | 9.0 | 128 | 29.4 | 40 | 15.9 | 2048 | - | - | [7–9] |
| hncacogpwg3d | 64 | 30% | 9.0 | 128 | 29.4 | 40 | 15.9 | 2048 | - | - | [9,12] |
| cbcaconhgp3d | 64 | 30% | 80.0 | 128 | 29.4 | 40 | 15.9 | 2048 | - | - | [11,13] |
| hncocacbgpwg3d | 16 | 25% | 80.0 | 128 | 30.0 | 40 | 15.9 | 2048 | - | - | [14] |
| hccconhgpwg3d3 | 64 | 25% | 80.0 | 128 | 35.0 | 40 | 15.9 | 2048 | - | - | [15–20] |
| noesyhsqcf3gpwg3d | 32 | - | 14.0 | 128 | 29.4 | 40 | 15.9 | 2048 | - | - | [21] |
| hnhagp3d | 32 | - | 29.4 | 40 | 14.0 | 128 | 15.9 | 2048 | - | - | [22,23] |
| hccconhgpwg3d2 | 64 | 25% | 15.9 | 128 | 35.0 | 40 | 15.9 | 2048 | - | - | [15–20] |
| noesyhsqcetgp3d | 32 | - | 14.0 | 128 | 80.0 | 64 | 14.0 | 2048 | - | - | [24] |
| hcchcogp3d | 16 | - | 14.0 | 128 | 80.0 | 64 | 14.0 | 2048 | - | - | [25] |
| hcchdigp3d | 16 | - | 14.0 | 128 | 80.0 | 64 | 14.0 | 2048 | - | - | [25] |
| hsqcnoesyhsqccngp4d | 8 | 30% | 80.0 | 32 | 10.0 | 64 | 35.0 | 32 | 15.9 | 2048 | [26] |
| hsqcnoesyhsqcccgp4d | 8 | - | 80.0 | 32 | 10.0 | 64 | 80.0 | 32 | 15.9 | 2048 | [11,21,27,28] |
| hsqcnoef3gpsi3d | 32 | - | 35.0 | 2 | 29.4 | 256 | 15.9 | 2048 | - | - | [21,29] |

**Supplementary Table 3.**

**Parameters used for calculations with ARIA 2.3.1 for the structure determination of TANGO1's cargo-recognition domain.**

| | |
|---|---|
| **General settings** | |
| Assignment frequency window | [ppm] |
| $^{1}H$ | 0.04 |
| $^{13}C/^{15}N$ | 0.5 |
| **Step 1 – Assignment and unrefined structure calculation** | |
| Violation tolerance [Å] | |
| Iteration 0 | 5.0 |
| Iteration 1 | 4.0 |
| Iteration 2 | 3.0 |
| Iteration 3 – 5 | 1.0 |
| Iteration 6 – 8 | 0.5 |
| Number of conformers | calculated/analyzed |
| Iteration 0 – 4 | 800/20 |
| Iteration 5 – 6 | 200/20 |
| Iteration 7 – 8 | 200/7 |
| Structures with no violations in iteration 8 | 33 |
| Simulated-annealing | |
| Distance restraint potential | log-harmonic |
| #steps high-temperature | 20,000 |
| #steps cooling 1 | 10,000 |
| #steps cooling 2 | 8,000 |
| **Step 2 – Structure refinement** | |
| Violation tolerance [Å] | |
| Iteration 0 | 0.5 |
| Number of conformers | calculated/analyzed |
| Iteration 0 | 6,000/50 |
| Refined structures | 200 |
| Structures with no violations after refinement | 76 |
| Simulated-annealing | |
| Distance restraint potential | flat-bottom |
| #steps high-temperature | 10,000 |
| #steps cooling 1 | 20,000 |
| #steps cooling 2 | 16,000 |

### Supplementary Figure 1

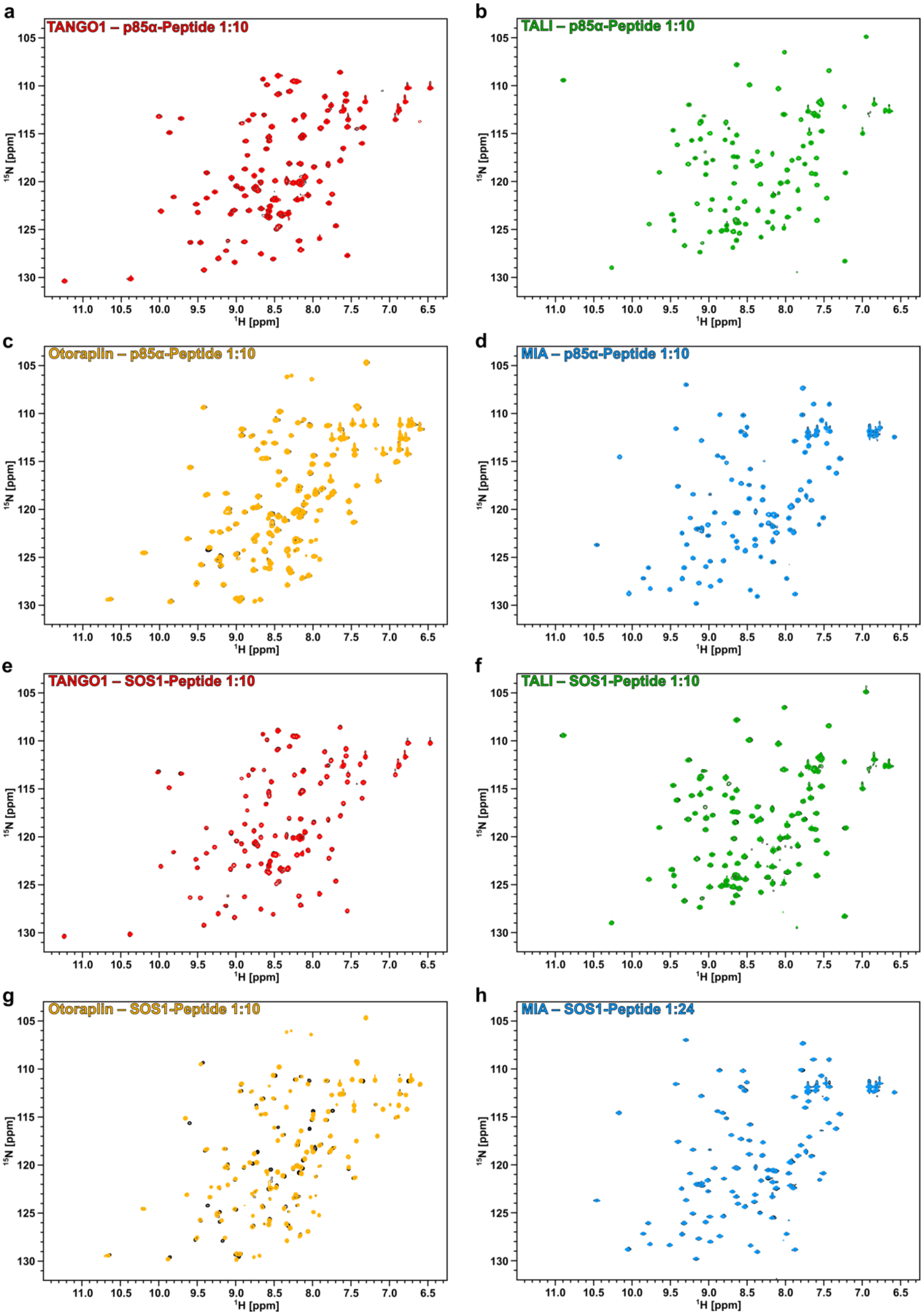

**$^{1}H^{15}N$-HSQC titration spectra of class I and II PPII ligands with homologous cargo-recognition domains encoded by the *mia* gene family.**

**a-d**, Titration spectra of MOTH domains from TANGO1, TALI, Otoraplin, and MIA, respectively, with an exemplary class I PPII ligand. The peptide was derived from residues 91 to 104 of the phosphatidylinositol-3-kinase regulatory subunit alpha (p85$\alpha$) known to interact with the SH3 domain of human tyrosine-kinase Fyn. [30] Reference spectra for all domains are shown in black. **e-h**, Analogous titration spectra with a class II PPII ligand. The peptide sequence corresponds to residues 1149 to 1158 of the guanidine exchange factor Son-of-Sevenless 1 (SOS1) of the Ras protein, reported to bind to the N-terminal SH3 domain of the growth factor receptor-bound protein 2 (Grb2). [31]

**Supplementary Figure 2**

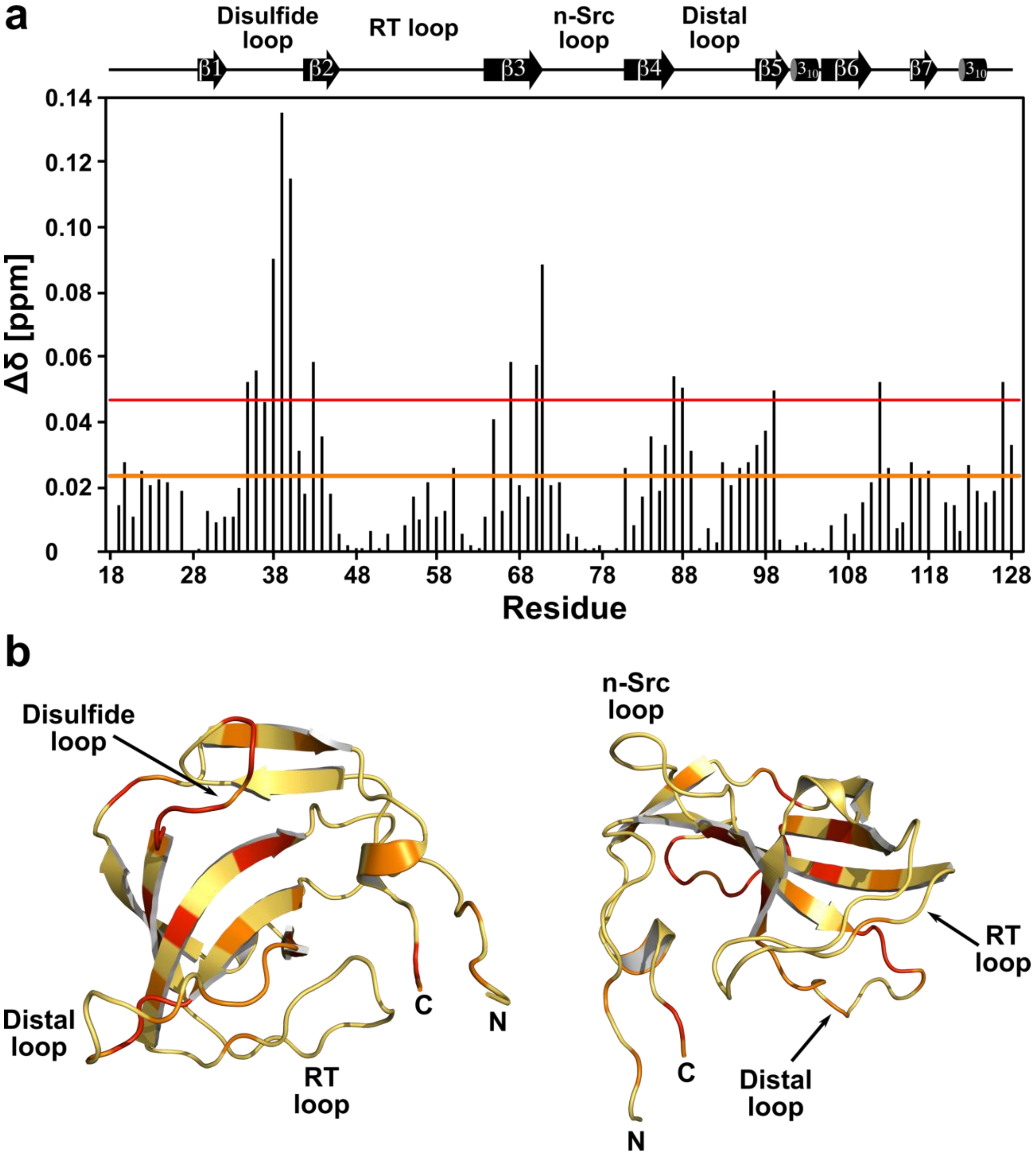


**CSP analysis of Otoraplin's interaction with a class II PPII ligand.**

**a**, Chemical shift differences between a reference spectrum and a tenfold molar excess of peptide plotted against the amino acid sequence. Orange and red line indicate the single and double standard deviation (SD), respectively, based on the average shift difference for all residues. **b**, Chemical shift differences exceeding the single (orange) or double (red) SD projected onto predicted structure of Otoraplin by AlphaFold (AF-Q9NRC9-F1. Accessed via https://alphafold.ebi.ac.uk/). Although Otoraplin exhibited a very weak interaction with class II ligands, and CSP mapping revealed the interaction site to be located opposite to the canonical SH3 binding groove between the RT and nSrc loop. Moreover, the interaction appears to be mainly mediated by electrostatic interactions between arginine side chains of the peptide and negative charges of the disulfide and distal loop of Otoraplin.

**Supplementary Figure 3.**

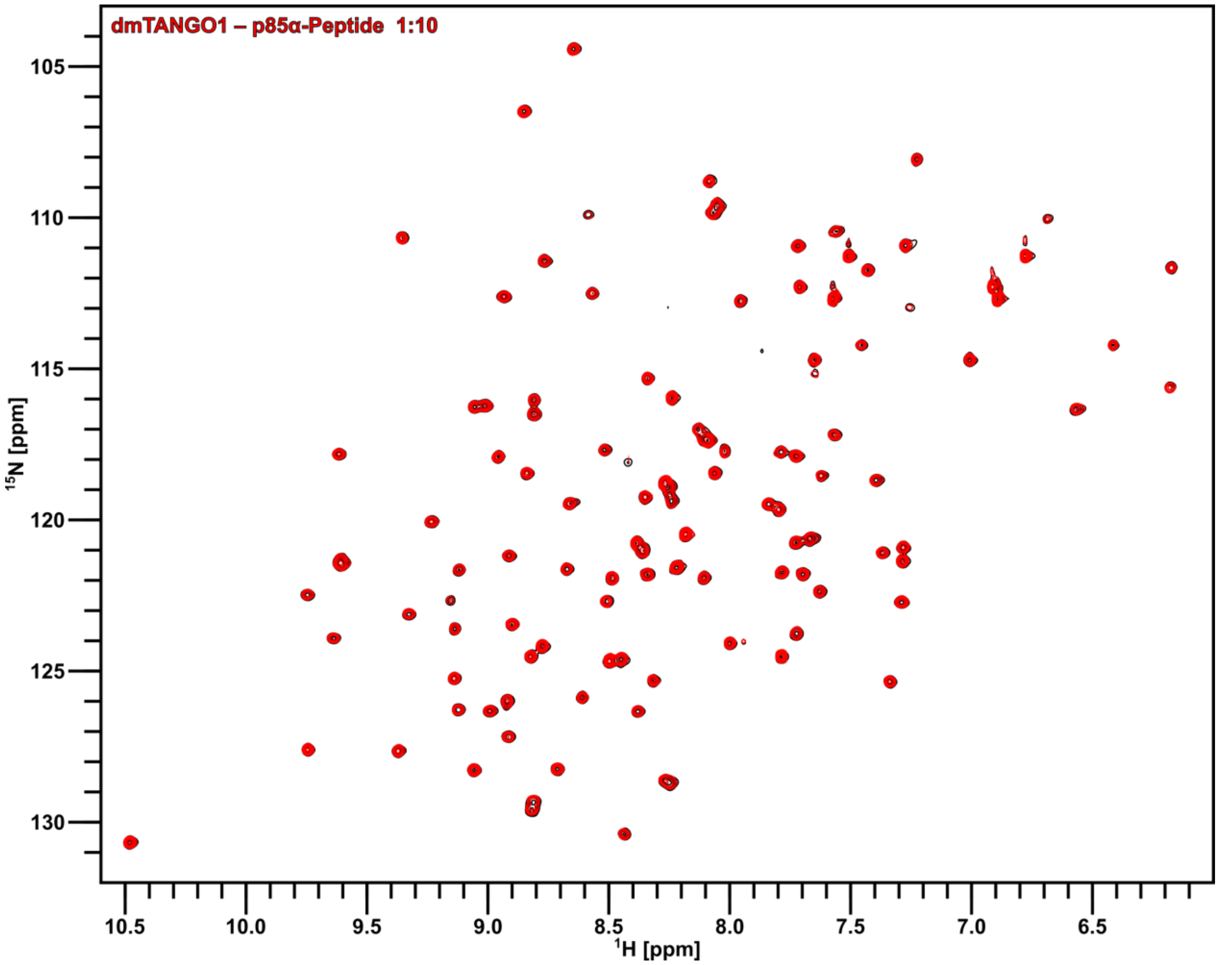


**$^{1}H^{15}N$-HSQC of dmTANGO1(30-139) and titration spectrum of PPII class I peptide.**

Titration spectra of TANGO1's cargo-recognition domain from *D. melanogaster* with an exemplary class I PPII ligand. The peptide was derived from residues 91 to 104 of the phosphatidylinositol-3-kinase regulatory subunit alpha (p85α) known to interact with the SH3 domain of human tyrosine-kinase Fyn. [30] Reference spectrum is shown in black.

Supplementary Figure 4.

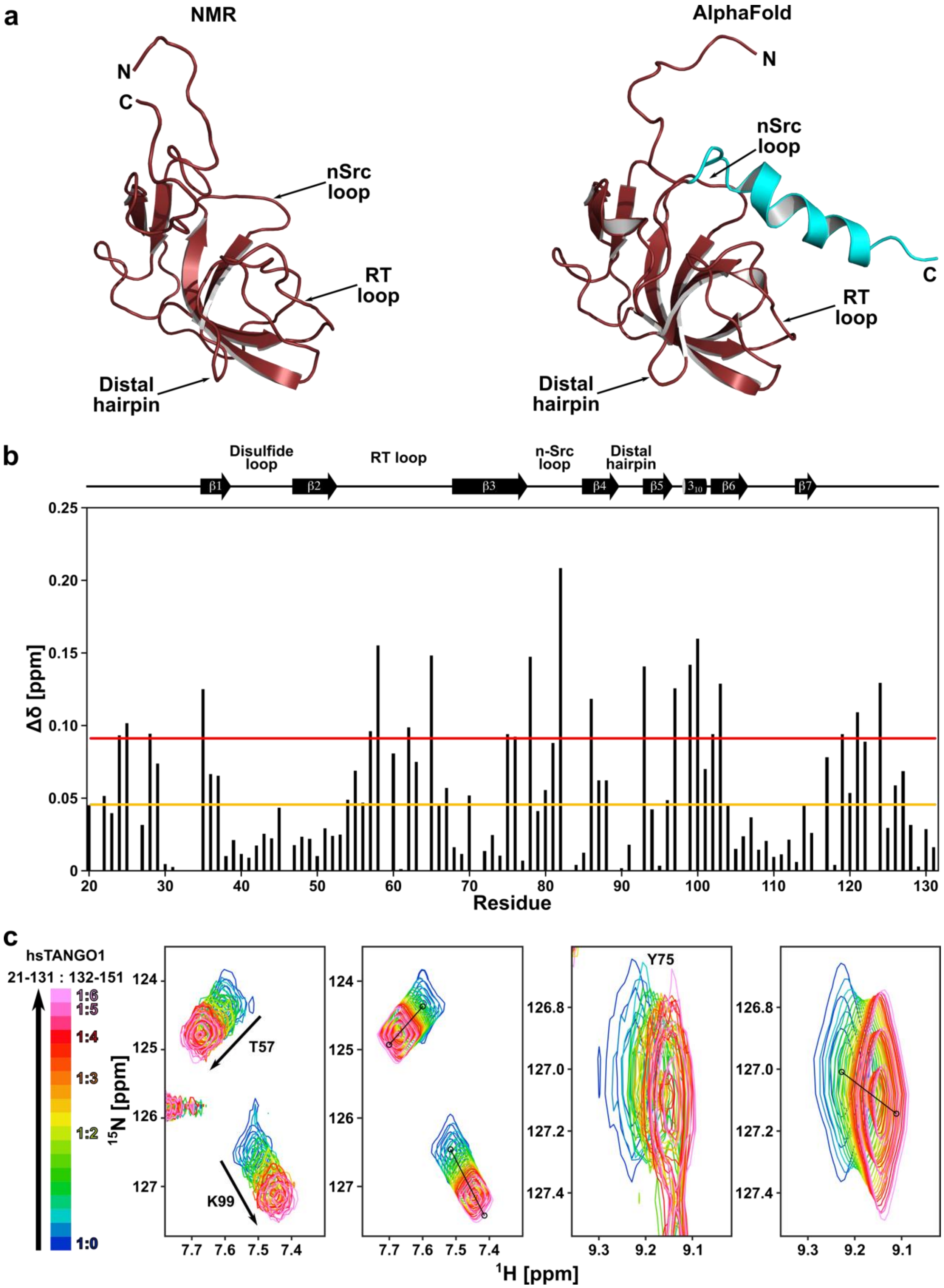

**Comparison of TANGO1's MOTH domain's structure from NMR spectroscopy and AlphaFold's structure prediction.**

**a**, Structures of TANGO1's MOTH domain from structure determination by solution NMR spectroscopy and prediction by AlphaFold display an RMSD for the structured region excluding the C-terminal helix of the prediction (cyan) of 0.84 Å. Most differences can be observed for non-secondary structure elements, predominantly for the nSrc loop and region between β-strands six and seven. These exhibit dynamic movements on the pico- to nanosecond timescale according to the hetNOE (Figure 2). **b**, Chemical shift differences observed for the human TANGO1 (21-131) upon titration of a peptide corresponding to residues 132-151 (displayed in **a** in cyan) with a sixfold molar excess. Single and double SD based on the average shift difference for all residues are indicated by orange and red line, respectively. **c** Exemplary excerpts of two-dimensional lineshape analysis of the interaction between human TANGO1(21-131) and a peptide corresponding to residues 132 to 151 of human TANGO1 using TITAN. Real spectra are displayed in respective left panels, re-calculated spectra based on fitted parameters are shown in the right panels.
